# Dynamic prioritization reshapes neural geometries for action in human working memory

**DOI:** 10.1101/2025.07.18.665374

**Authors:** Ivan Padezhki, Juan Linde-Domingo, Carlos González-García

## Abstract

Prior work has shown that control brain regions encode upcoming novel instructed actions. Similarly, visual working memory (WM) representations can reflect different priority states. However, it remains unknown whether the priority status of WM-guided novel actions similarly modulates their neural coding format, and how such dynamics unfold over time. We addressed these questions using EEG while human participants performed two consecutive choice reaction tasks. At the start of each trial, participants encoded two novel stimulus-response (S-R) mappings. A cue then indicated whether these mappings would be relevant immediately (“current” condition), later after an intervening task (“prospective”), or after a free delay (“delayed”). Using multivariate pattern analysis, we found that the S-R category was decodable in all conditions during relevant time windows. Critically, pattern similarity analyses revealed that while mere maintenance demands allow for temporary preservation of neural codes (i.e., between current and delayed trials), shielding from interference (i.e., prospective trials) induced significant alterations to the neural code. Further exploration of the representational geometry revealed that priority status gained prominence dynamically over S-R category coding when preparing for such shielding demands. Importantly, some of these changes emerged anticipatorily, prior to target onset. Overall, our results show that, similar to visual WM, the priority of intended actions dynamically and anticipatorily reshapes their neural format. They further reveal how different demands induce content geometries compatible with previously proposed coding schemes, and that such representational changes can be implemented flexibly in time.

**Significance statement:** Successfully translating visuomotor plans into action relies not only on maintaining such plans but also on prioritizing relevant information at the right time. While prior work has shown that working memory representations adapt to prioritize visual content, it remains unknown whether similar mechanisms apply to action-guiding content. Using EEG and multivariate analyses, we show that priority states—whether immediate, delayed, or prospectively relevant—modulate the representational geometry of novel stimulus-response mappings. Notably, the anticipation of interference compared to mere maintenance leads to greater alterations in the neural code during a retention interval. These results demonstrate that neural representations of novel instructed actions are shaped by their priority state and upcoming demands and reveal how goal- relevant information is restructured over time.

## INTRODUCTION

To reach our goals, we frequently have to maintain and manipulate information no longer present as sensory input. The set of computations that allow such ability is usually referred to as working memory (WM). WM serves as a cornerstone for adaptive behavior and is fundamental in a wide range of higher-level functions (Baddeley, 1992; Oberauer, 2009).

Significant effort has been devoted to describing the cognitive and neural architecture that might support WM, with special emphasis on aspects such as its capacity limits or its temporal constraints (Cowan, 2010; Formica et al., 2024; Fukuda et al., 2010; Shepherdson et al., 2018). Recently, another feature of WM has gained prominence in the field: its flexibility, or how stored information can be dynamically prioritized based on its relevance. Theoretical frameworks and experimental evidence suggest that internal endogenous attention acts as a priority switch (Myers et al., 2017; Nobre & Stokes, 2020) and that different states of priority are represented dynamically in visual WM (Muhle-Karbe et al., 2021; Myers et al., 2018; Stokes et al., 2020; van Ede, 2020).

While prioritization of visual information has been widely studied (de Vries et al., 2018; Linde-Domingo & Spitzer, 2024; Olmos-Solis et al., 2021; van Loon et al., 2018; Wan et al., 2022; Yu et al., 2020), less is known about more complex WM contents, such as planned actions or stimulus-response (S-R) associations, where enabling an optimal coding format is essential for adaptive behavior (González-García et al., 2021). Think of the association between a sink and washing your hands. Sometimes, if you are washing your hands before eating, you might need to keep in WM the action plan to perform it *immediately*, while other times, you might have to do something else before. These scenarios show that S-R associations seem to be held in WM under different priority states. Recent behavioral evidence suggests that planned actions can be dynamically prioritized as required by the task at hand and, in turn, that their functional state influences ongoing behavior (Formica et al., 2024).

A critical question is how planned actions are maintained in neural patterns under different states of priority and how they interact with each other. In visual WM, recent theoretical proposals have put forward three main possibilities for such an interaction (Stokes et al., 2020). First, attentional gain coding proposes the involvement of the same neural populations in the two representations but with different degrees of activation, predicting a positive correlation between them. Second, suppressive coding expects the involvement of the same subpopulations, but with suppressed activation for prospective contents. This scheme is thought to shield prospective memories from interference and predicts a negative correlation between patterns. Finally, orthogonal coding predicts no correlation between the patterns. While there exists some evidence in visual WM, it remains unknown whether the state of priority of WM-guided novel actions similarly modulates their representational format. Moreover, the temporal unfolding of such effects is unclear.

Here, we used electroencephalography (EEG) while participants encoded two novel S-R mappings at the beginning of each trial. Subsequently, a cue indicated whether the encoded mappings would be relevant in a first probe (“*current*” condition), in a second probe after having to answer to an irrelevant stimulus (“*prospective*” condition), or in the second probe after a free delay (“*delayed*” condition). As such, the current versus delayed comparison informs about differences attributable to pure maintenance demands, while comparing delayed to prospective captures differences driven by the need to shield relevant S-R mappings from interference. Lastly, the current versus prospective comparison emphasizes differences due to both maintenance and shielding demands. We then used Representational Similarity Analyses (Kriegeskorte et al., 2008) to explore how patterns from different conditions related to each other. Our results suggest that the priority state of WM-guided novel actions proactively impacts their representation. They also provide initial evidence that the effect of priority is dynamic, modulating the representation of WM contents throughout the entire time they are maintained.

## METHODS

### Participants

Thirty-four healthy participants (mean age = 22.15 years, range = 18-30; 23 females, 11 males) recruited from the participant pool from the University of Granada took part and received a compensation of 10 euros per hour. Participants either had normal vision or wore glasses that matched their correction. The study was approved by the University of Granada’s Ethics Committee, and all participants provided informed consent before starting the experiment. A total of 3 participants were excluded from all analyses due to their error rate being above 60% in probes 1 and 2 in regular trials (Formica et al., 2024) leaving 31 participants for further analyses.

### Apparatus, stimuli, and procedure

S-R associations were formed by pairing images with their screen position and words that indicated required response fingers (either “índice” or “corazón”, Spanish words for “index” and “middle”, respectively). On each encoding screen (see Fig. 1A), one of these words appeared centrally, surrounded by two images (one on the left-hand side of the screen and another on the right-hand side of the screen). While the word indicated the response finger, the position of the image indicated the specific effector. For instance, an encoding screen displaying a fox on the left-hand side, an elephant on the right-hand side, and the word “índice” centrally, would specify two mappings: fox-left index finger and elephant-right index finger. This allowed us to have 2 mappings on screen that involved the same (stimulus and) response category (e.g. index finger) but different effectors (e.g. left index finger vs right index finger).

**Figure 1.**
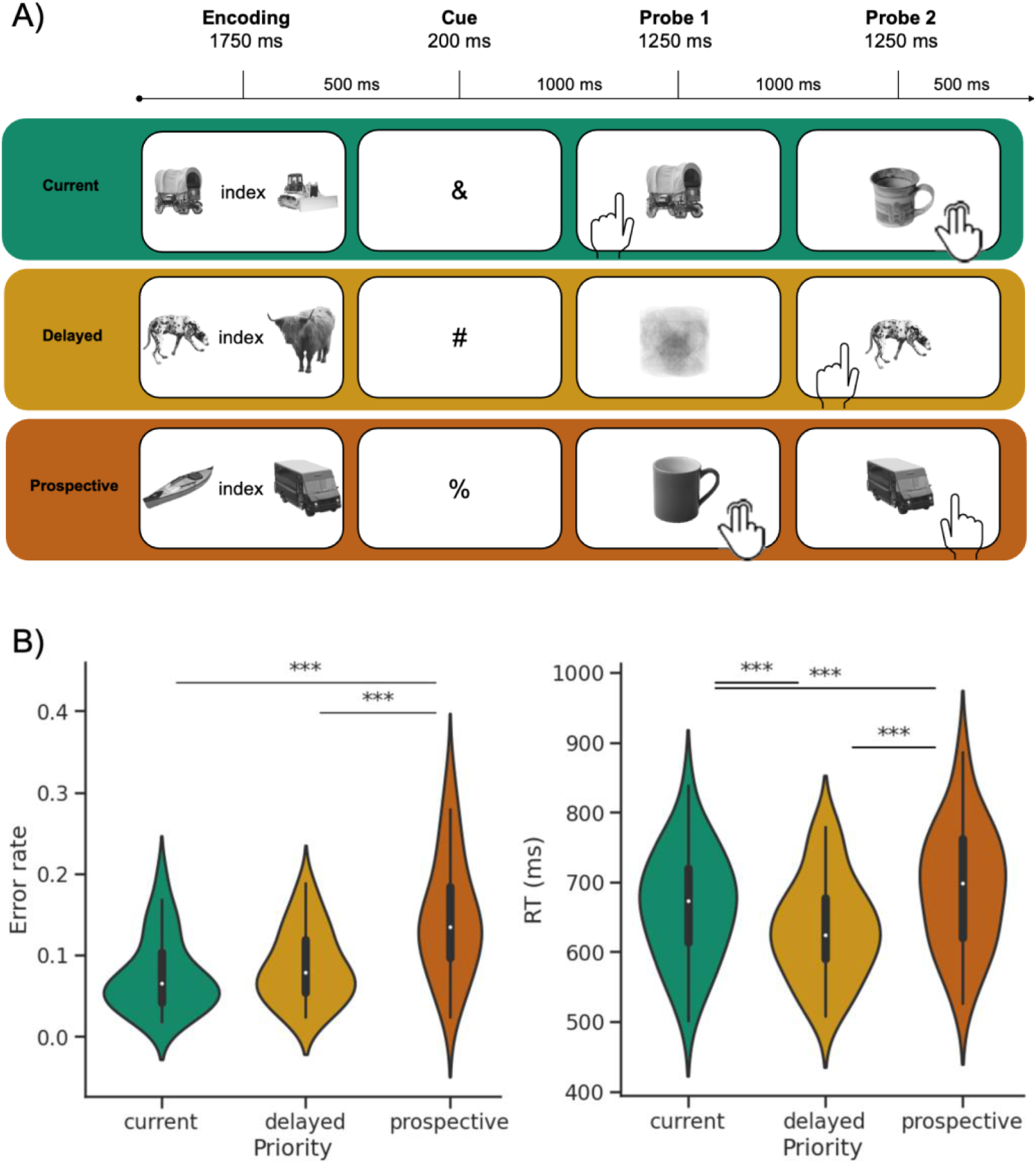
A) Experimental paradigm: On each trial, participants first encoded two novel S-R mappings consisting of the association between an item (mammal or vehicle) and a response. After the encoding screen, a cue indicated when in the trial the encoded mappings and an additional irrelevant mapping would be relevant (relevant in the first probe, relevant in the second probe after an irrelevant task, or tested in the second probe and no task required in probe 1). Then, two consecutive target stimuli prompted participants to provide the associated response (either the instructed response, for relevant mappings, or the overlearned response for irrelevant mappings). B) Error rate (left) and reaction time (right) behavioral results. Stars denote significant post-hoc tests (p < 0.05 after correcting for multiple comparisons).

Each relevant S-R association was presented just once during the entire experiment to prevent the formation of long-term memory traces (Beukers et al., 2021; Meiran et al., 2015). Given this prerequisite, images of mammals and vehicles were compiled from different available databases (Brady et al., 2008, 2013; Brodeur et al., 2014; Griffin et al., 2006; Konkle et al., 2010), creating a pool of 821 unique pictures (431 mammals, 390 vehicles). To increase perceptual similarity and facilitate recognition, the background was removed from all images, items were centered on the canvas, and they were converted to greyscale images.

Importantly, even though specific S-R associations were presented only once throughout the experiment, they could be grouped depending on the specific combination of stimulus and response dimensions. Specifically, the combination of the 2 stimulus dimensions (mammals/vehicles) and the 2 response dimensions (index/middle finger) leads to 4 finger-animacy pairings: S-R 1 (mammals-index), S-R 2 (mammals-middle), S-R 3 (vehicles-index), and S-R 4 (vehicles-middle).

On each trial, participants were presented with two consecutive probes on which they had to perform two choice reaction tasks, i.e., a task involving the implementation of condition-action rules (Barrouillet et al., 2015; Gade et al., 2014; Oberauer, 2010), except in the Delayed condition (see below). Each trial started with an encoding screen (1750 ms) that displayed 2 S-R associations as described above. After a 500 ms fixation period, a symbolic cue indicated when, in the trial, the encoded mappings would be relevant. For instance, in some blocks, a “&” cue indicated that the encoded mappings were relevant for the first choice reaction task and irrelevant for the second, where an irrelevant mapping would be probed (see next paragraph for details on irrelevant mappings). This constituted the Current condition. The Prospective condition, in contrast, was signaled with a “#” cue and instructed participants to use the encoded mappings in the second probe and to expect to use an irrelevant mapping in the first probe. Last, the Delayed condition was cued with a “%”, instructing participants to wait for the second probe to execute one of the encoded mappings, and no task was required during the first probe. Symbolic cues were counterbalanced across blocks so that at the end of the experiment, all of them had been equally associated with all three conditions. To reduce the cost of switching from one cue meaning to another, blocks using the same symbol assignment were grouped.

Regarding irrelevant mappings, to avoid extra WM load and a longer practice session, we used objects that automatically cue left vs. right-hand actions (i.e., mugs with the handle on the right- or left-hand side), and participants were asked to press the two keys of the hand towards which the handle was pointing. To reduce the predictability of the irrelevant mappings, we used 30 different mug exemplars. We also created filler images for the Delayed condition (see below) by averaging 25 objects not used in any other probe.

In all cases, the cue was followed by a fixation point (cue-target interval; CTI) for 1000 ms. Directly after the CTI, the first target was on screen for 1250 ms (see Fig. 1A). In the Current condition, the first target screen displayed the stimulus of one of the two encoded mappings, prompting participants to execute the associated response by pressing the corresponding key. In the Prospective condition, the first target consisted of a picture of a mug, prompting participants to execute the irrelevant mapping as described above. Last, in the Delayed condition, a filler image was displayed for the same duration (1250 ms), and participants were instructed to simply wait for the second probe. After Probe 1, a fixation point was shown on the screen (inter-stimulus interval, ISI) for 1000 ms, before the display of a second probe for 1500 ms. Here, target screens followed the same logic as the first probe, but in this case, a relevant stimulus was shown in the Prospective and Delayed conditions, and an irrelevant one in the Current condition. Last, a fixation point was shown between trials (inter-trial interval, ITI) for 500 ms. Additionally, in ∼6% of relevant targets, a catch image appeared. This consisted of a new image, different from any of the encoded stimuli, to which participants had to answer by pressing the 4 available buttons on the keyboard. Catch trials were included to ensure that participants encoded the two S-R associations on every trial and were randomly distributed across priority conditions. Overall, each trial lasted on average 7 seconds. The sequence of trial events is depicted in Figure 1.

The main task was divided into 9 blocks. Each contained 60 trials (55 regular and 5 catch trials), and 20 per priority condition (current, prospective, delayed). The four possible S-R pairings were fully counterbalanced across priority conditions, resulting in 5 trials per crossing of priority and S-R on each run and 45 across the entire experiment. Each run lasted around 8 minutes, and the main task, containing 540 trials, lasted around 70 minutes in total. Before the main task, participants performed a practice session with trials following the same structure described above, except that feedback was included to help familiarization.

### Behavioral and EEG data acquisition

Behavioral response recording and stimulus presentation were performed using PsychoPy 2 (Peirce, 2007) running on a Windows PC and projected onto a screen. Participants responded using the keyboard buttons s, d (left hand), and k, l (right hand) on which participants placed their index and middle fingers. The EEG data were collected in a Faraday room using 64 active electrodes (actiCap, Brain Products, GmbH, Munich, Germany) that were positioned following the international 10% system. During collection, almost all electrodes showed an impedance lower than 10 kΩ except for some electrodes (Fp1, Fp2, AF7, and AF8), which were noisier for all subjects. Data were recorded using a BrainAmp DC amplifier (Brain Products, GmbH, Munich, Germany) at a sampling rate of 500 Hz sampling rate.

### Behavioral data pre-processing

Besides the exclusion of three subjects based on their low accuracy in the two probes, no other steps were applied for the behavioral analyses. After taking the relevant trials, for each of the three conditions, an average of 165.00 (±2.70) trials remained for behavioral analyses.

### EEG Pre-processing

For the EEG analyses (carried out in the MNE Python package, version 1.8.0), further exclusion criteria were applied to ensure good data quality. Firstly, the data for each subject was inspected visually, and muscular activity artefacts were marked for removal. Noisy channels were identified by inspecting the raw data. If subjects had less than 15% noisy channels, those channels were interpolated using the spherical spline method, as explained in Perrin et al. (1989). Otherwise, the subjects would have been excluded if they had more than 15% noisy channels, but we did not observe this in the sample. In this method, electrode locations are projected onto a unit sphere, and neighbor channels are used to interpolate noisy ones. Blink and eye movement components were identified and removed from the data using independent component analysis (ICA). This involved removing one component for blinks and one for eye movements for all subjects. In the next step, common values were set for a band-pass filter (0.5 Hz to 60 Hz) along with a band-stop filter (48–52 Hz) to remove line noise.

After merging the trial information with the behavioral output file, trials were formed. An epoch spans all experimental events in a trial, covering 8 seconds. Based on two further selection criteria, certain trials were excluded. For one, the trials that included muscular noise (previously annotated on the raw data through visual inspection) were excluded. Additionally, trials with slow drift (peak-to-peak channel amplitude) were excluded (threshold: 150 μV). Based on these two criteria, from the original 540 trials per subject, an average of 429.58 (± 97.94) trials would remain; however, the range (110, 526) suggested that some subjects lose a significant number of trials in this step. For this reason, three subjects who lost more than 50% of their trials were excluded from further analyses. This decision was further motivated by the type of analyses, which requires a minimum amount of trial to ensure stable RDM estimation for each subject, along with reliable cross-subject comparison of RDMs (Grootswagers et al., 2017; Walther et al., 2016).

The data was re-referenced using the average reference method. Epochs of 8 seconds were extracted, starting from 0.5 seconds before encoding screen onset to 7.5 seconds afterwards, covering all experimental events in the trials. Another exclusion was performed based on the subjects’ success in the catch trials, namely, participants were excluded if they answered these using a single key press (that is, as if it were a regular trial) in more than 50% of cases. Therefore, 5 subjects that fell below this threshold were excluded, resulting in a final sample of 23 subjects for the EEG analyses. Finally, only non-catch trials and correct trials were selected for further analysis. This left a range between 296 and 446 trials per participant remaining (Mean: 382.13, Std: 41.07). Baseline correction was applied using the mean signal from -0.5 to -0.1 seconds relative to encoding onset. For all analyses, the data were downsampled to 50 Hz given the length of the epochs, that is, full trials.

### Representational Similarity Analysis

To prepare each subject’s data for RSA, demeaning was applied across all epochs. Then, trials were ordered by event type: animacy and response finger. The data from the three WM conditions were then concatenated to form a 12-by-12 matrix with 3 conditions and 4 categories in each. Then, the data was split into three parts for a 3- fold cross-validation. Within each fold, a whitener was trained on the training set, consisting of 2 partitions, and applied to the training and test sets, where the test set is the third data partition. The whitening process is a multivariate noise normalization method (Guggenmos et al., 2018) which normalizes each channel by estimating its covariance matrix. The goal is to emphasize sensors with low noise while downweighing noisy ones. (In detail, the process includes computing the covariance matrix - SVD with a shrinkage parameter of 1e-5 - for each condition across sensors. The whitening matrix is the inverse square root of the covariance matrix. It is post-multiplied with the subject’s data.) The cross-validation scheme was as follows: within a fold, two out of three partitions of the trials were taken for the training set and averaged, while the other partition formed the test set. The distance between the train and test sets was computed and later averaged over the distances from the 3 folds. The cross-validated representational dissimilarity matrix was calculated as 1 - Pearson correlation. For analyses using similarity (instead of dissimilarity) matrices, we subtracted the RDM from 1 to get the correlation.

Depending on the metric calculated, we took each subject’s RDMs over the time course of the trial and averaged these over the whole sample. In other words, an RDM was cross-validated independently for each time point during the trials of each subject.

### Cluster-based permutation test

A cluster-based permutation test was performed to assess the significance of the metrics calculated. The MNE function for a 1-sample permutation cluster test uses sign flips to randomize the data. For the RSA analyses, a two-tailed test was performed with 2048 permutations. The cluster-forming threshold was calculated using a p-value of 0.01 to get a corresponding distribution of t-values. Then, clusters with a p-value below 0.05 and fewer than 15 consecutive time points were discarded.

## RESULTS

### Task priority modulates behavioral performance

We first analyzed behavioral outcomes during relevant probes. A repeated-measures one-way ANOVA yielded strong evidence for a main effect of priority conditions on reaction times (RT: F = 58.25, p < 0.001, ηg² = 0.086). Post-hoc tests revealed faster RT in Delayed trials (*M* = 633.6ms, *SD* = 72.4ms) compared to Prospective trials (*M* = 692.5ms, *SD* = 86.9; t = -10.44, p < 0.001, BF10 = 5.52e+08) and Current trials (*M* = 666.0ms, *SD* = 78.7; t = 6.211, p < 0.001, BF10 = 22280.0), as well as faster RT in Current than in Prospective trials (t = -4.796, p =, BF10 = 573.167). We also found strong evidence for a main effect of Priority on error rates (ER: F = 46.86, p < 0.001, ηg² = 0.244). As with RT, participants made fewer errors in Delayed trials (*M* = 0.087, *SD* = 0.045; t = -7.485, p <0.001, BF10 = 566.70) than in Prospective (*M* = 0.148, *SD* = 0.073). Similarly, they made fewer errors in Current (*M* = 0.076, *SD* = 0.046; t = - 7.30, p <0.001, BF10 = 352.90) than in Prospective trials. In contrast, the evidence was inconclusive for the difference between the Current and Delayed trials (BF10 = 1.184).

### Representation of relevant S-R mappings under different priority demands

We first investigated whether the relevant stimulus-response mapping of each trial was reflected in EEG patterns. To do so, we computed a Representational Dissimilarity Matrix (RDM; see methods) for each time point of the epoch, across trials within the same priority condition. Using the obtained RDMs, we then obtained a S-R Discrimination Index (SRDI) by averaging the within-condition lower triangles of each RDM and subtracting the mean within-condition diagonal from such average (see Fig. 2A). The SRDI thus provides an estimate of whether, at a given time point, EEG activity patterns discriminate between the relevant S-R mapping among the four S-R categories. Results revealed significant clusters of SRDI values during sensitive trial events (all p-values < 0.01; see Fig. 2B). Specifically, in Current conditions trials, relevant S-R mappings were discriminable during the presentation of the first [3.92s to 4.24s relative to encoding onset, peak at 3.98s with a value of 0.09] but not the second probe, whereas in Delayed and Prospective trials, the opposite pattern was observed [Current condition, 3.80s to 4.36s relative to encoding onset, peak at 4.00s with value 0.08; Delayed, 6.08s to 6.62s relative to encoding onset, peak at 6.18s with a value of 0.12; Prospective, 6.02 to 6.30s relative to encoding onset, peak at 6.20s with a value of 0.10]. Interestingly, in contrast with those times related to the relevant probe, no significant SRDI clusters were found during the cue-target interval, suggesting no evidence of preparatory reactivation of category information, regardless of the priority condition. Bayesian statistics provided evidence in line with frequentist tests. Altogether, these results indicate that the specific S-R mapping of each trial was reflected in the EEG signature. Still, such activity was limited to the relevant probe of each condition.

**Fig 2.**
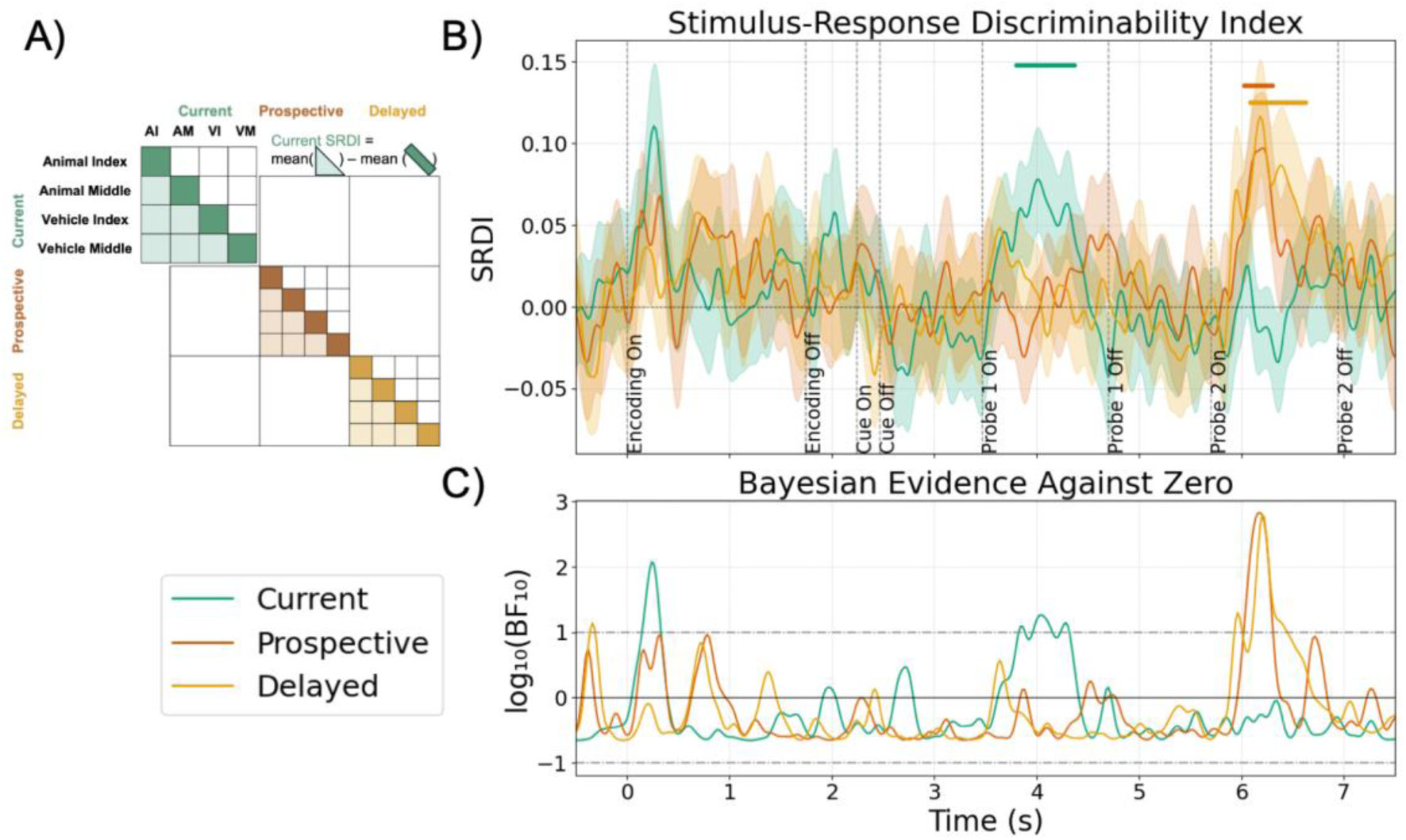
**S-R Discriminability Index per Priority Condition**. A) Analysis rationale. B) SRDI time-resolved results. Color straight lines denote significant SRDI values above 0 (p < 0.01, cluster-based corrected for multiple comparisons). C) Bayesian evidence for results in panel B. Horizontal dashed lines at 1 and -1 denote strong evidence for the alternative (SRDI different from 0) and the null hypothesis (SRDI = 0), respectively. Values between ±0.5 and ±1 reflect moderate evidence, while values between 0.5 and -0.5 reflect inconclusive evidence.

**Fig 3.**
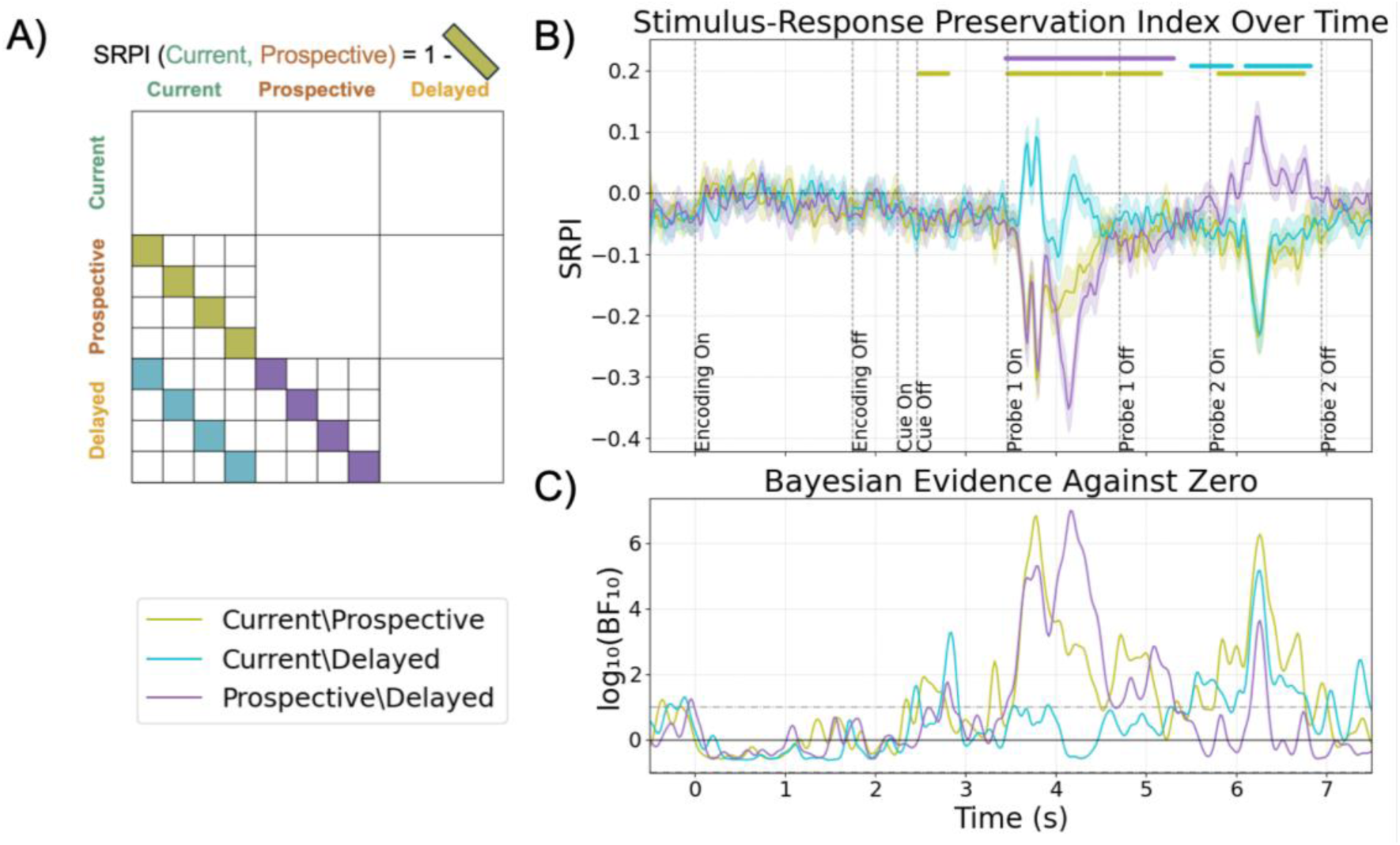
**S-R Preservation Index between Priority conditions**. A) Analysis rationale. B) Preservation of time-resolved results. Color straight lines denote significant SRDI values above or below 0 (two-sided t-test against 0, p < 0.01, cluster-based corrected for multiple comparisons). C) Bayesian evidence for results in panel B, using the same evidence thresholds described in Fig. 2D.

### Priority demands impact the preservation of neural codes

The within-category results in the previous section revealed widespread S-R discriminability during relevant trial probes. However, these results are agnostic about the similarity of S-R neural codes under different priority conditions. To investigate whether S-R mappings are represented similarly across Current, Delayed, and Prospective trials on EEG patterns, we derived an S-R Preservation Index. This pattern was obtained by averaging the lower triangle of the between-condition diagonals of Representational Similarity Matrices from each time point (see Methods for details). In this analysis, a Preservation value larger than 0 suggests a generalizable or common code between conditions (i.e., positively correlated neural pattern), whereas values below 0 can be indicative of negatively correlated neural patterns (i.e., suppressive coding). Finally, a preservation of 0 suggests no relation between the codes in the two conditions (i.e., orthogonal codes).

Our results revealed two negative Preservation clusters during the first probe when comparing Current and Prospective [3.46s to 4.50s relative to encoding onset, peaking at 3.78s with value -0.31, directly followed by a cluster from 4.56 to 5.16s, peaking at 4.90s with -0.11], as well as when comparing Prospective and Delayed trials [3.44 to 5.30s relative to encoding onset, peaking at 4.14s with value -0.35]. This suggests that, in Probe 1, the patterns of neural activity in Prospective trials are consistently anticorrelated with those of the other conditions. However, this organisation of the neural activity changed during Probe 2, where the code between Current trials and the other conditions was found to be significantly anticorrelated: the S-R Preservation index for Current versus Prospective trials shows a significant negative cluster between 5.80 and 6.74s with a peak of -0.24 at 6.24s. Similar results are observed for Current versus Delayed trials during Probe 2: two subsequent clusters, starting from 5.50s to 5.94s with a peak of -0.08 at 5.74s, followed by a cluster from 6.10s to 6.82s peaking at 6.26s with a value of -0.23. Importantly, this analysis revealed a significant cluster in the cue-target interval when comparing Current and Prospective (from 2.48s to 2.80s, absolute peak at 2.52s with a value of -0.06), suggesting that the suppressive coding pattern between these two conditions observed during the first probe takes place in a preparatory manner, after the retro-cue is presented. Additionally, Bayesian tests provided evidence for preparatory preservation activity between the Current and Delayed conditions (from 2.72 to 2.92 s). These results reveal an intricate relationship where the code of Current and Delayed trials is anticorrelated during the cue-target interval, but becomes positively correlated after the cue of the first probe, suggesting again a dynamic reconfiguration of neural patterns as a function of task demands and trial events.

### Priority demands dynamically affect the geometry of S-R representations

To further investigate which factors influence the neural geometry in our task, we derived a Condition-Relative Discriminability Index (CRDI). This index consists of the subtraction of two subparts of RDMs: first, the distances between the 4 S-Rs in one condition; second, the distances between the S-Rs in this same condition and the rest of the conditions. Intuitively, CRDI describes the relative influence of the priority condition vs. the S-R category on the neural geometry at a specific time point. Specifically, when negative (below zero values), it reflects a greater contribution of the priority condition to the underlying neural patterns, whereas above zero values reflect a greater role of S-R categories.

Similar to the metrics reported before, results reveal a dynamic effect of priority demands across trial events. Interestingly, CRDI values in the Current condition became negative during the second probe [5.94s to 6.50s relative to encoding onset, peaking at 6.26s with a value of -0.23], reflecting that representations in Current trials in this epoch are largely dominated by the priority condition. In Prospective trials, this pattern was reversed. During the first probe, we observed negative CRDI values [3.54s to 3.84s relative to encoding onset, peak of -0.26 at 3.80s]. For Delayed trials, positive CRDI values were found during the second probe [6.04s to 6.42s relative to encoding onset, peaking at 6.26s with a value of 0.14].

When taking a look at CRDI values from a Bayesian perspective (see Fig. 4c), we found evidence supporting a similar pattern for Current and Prospective trials: for their relevant probes CRDI values are positive (Current trials, log₁₀(BF₁₀) = 2.67 from 3.64s to 3.74s relative to encoding onset, followed by another cluster from 4.08s to 4.30s, with a value of 3.23; Prospective trials, log₁₀(BF₁₀) = 1.67 from 6.14s to 6.20s relative to encoding onset) but negative in irrelevant probes. (Current trials, log₁₀(BF₁₀) = 6.58 from 5.96s to 6.50s relative to encoding onset, followed by another cluster from 6.60s to 6.80s with log₁₀(BF₁₀) = 4.54; Prospective trials, log₁₀(BF₁₀) = 6.63 from 3.58s to 3.84s relative to encoding onset, and a subsequent cluster from 4.12s to 4.30s with log₁₀(BF₁₀) = 2.33). Interestingly, in Delayed trials, Bayesian analyses revealed evidence for positive CDRI values during both Probe 1 (from 3.74s to 3.82s with log₁₀(BF₁₀) = 3.09) and Probe 2 (from 6.06s to 6.40s with log₁₀(BF₁₀) = 2.27).

**Fig 4.**
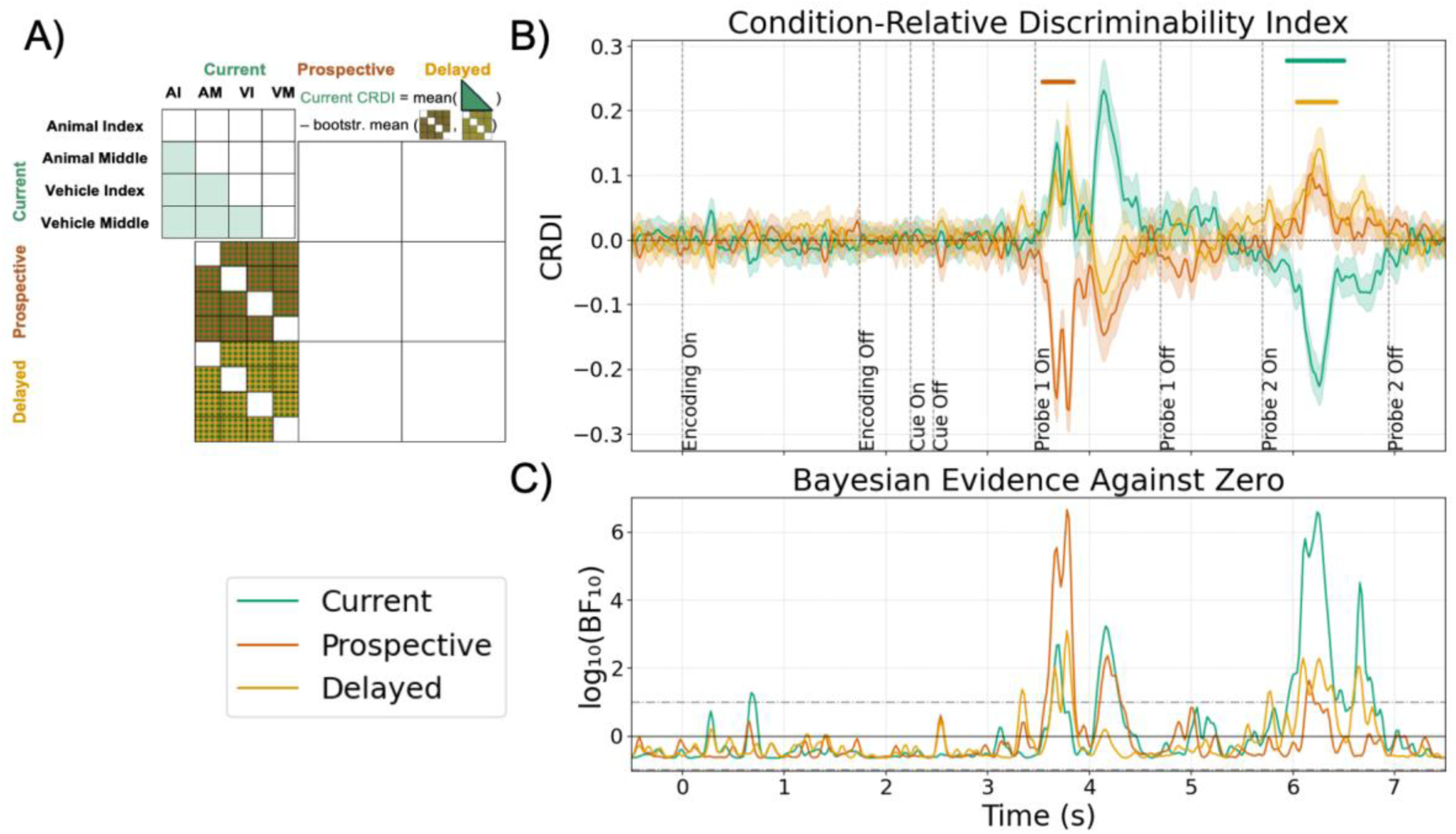
Condition-Relative Discriminability Index of S-Rs. A) Analysis rationale. Unlike the construction of the previous indices, here we consider the interplay between S-R mapping and the Priority condition in determining the strength of the similarity. The index is computed using the distances among the four S-R associations within one WM condition and the distances among this condition’s S-Rs versus the other conditions. As the lower triangle sub-matrices are not symmetrical along their diagonals, both sides were used in the computation of the index. In addition to this, we computed the Monte Carlo bootstrapped mean of these cells to match the sample size of the within-condition lower diagonal, which was estimated using a larger number of trials. B) Time-resolved results. Colour straight lines denote significant CRDI values above or below 0 (two-sided t-test against 0, p < 0.01, cluster-based corrected for multiple comparisons). C) Bayesian evidence for results in panel B, using the same evidence thresholds described in Fig. 2D.

Altogether, these results indicate that the neural geometry underlying WM retention periods dynamically adapts to task demands and can prioritize or minimize the contribution of different dimensions.

## DISCUSSION

In the current study, we aimed to understand how the prioritization of novel S-R mappings in WM altered their neural representation. By combining EEG with multivariate pattern analyses, we show that not only are novel action plans reliably detected in EEG activity during relevant trial epochs, but also that their neural format is substantially shaped by their priority status. Specifically, while mere maintenance demands (i.e., delayed condition) had limited impact on the neural patterns, shielding relevant S-R associations from interference (i.e., interference condition) led to marked changes in representational geometry. These changes were reflected in reduced similarity with the other conditions and shifts in the relative contribution of task-relevant dimensions. Importantly, some of these geometrical motifs were observed prior to target onset, revealing an anticipatory component. Altogether, these results suggest that the format of action-guiding representations in WM is not static but instead dynamically adapted based on their immediate or future relevance.

Previous research has shown neural priority effects on visual WM, where representations of currently relevant items differ from those that are prospectively relevant (Stokes et al., 2020). Recently, much emphasis has been put on how the format of neural codes might differ between different priority demands. Critically, previous studies have offered mixed results in this regard. While some findings suggest shared but attenuated codes for prospectively relevant items (in support of attentional gain models; Chun, 2011; Schneegans & Bays, 2017), others point to orthogonalization or even inversion of neural patterns for latent representations (van Loon et al., 2018; Yu et al., 2020). Moreover, these previous studies have focused on visual contents, but evidence for more complex, action-related WM representations is only recently growing. First, there is behavioral work demonstrating that novel instructed actions, such as simple S-R mappings, can be flexibly favored and that this prioritization influences task performance (Formica et al., 2024). Moreover, neuroimaging studies suggest that the neural patterns supporting the implementation of novel instructed tasks are dynamically organized within a trial (Pena et al., 2025). Our study bridges these two strands of research by providing a direct characterization of the neural representations of S-R associations across multiple levels of priority.

Our results suggest that different priority demands induce different coding schemes. We obtained evidence for a suppressive coding model governing the relationship between currently and prospectively relevant S-R associations, as revealed by the systematic anticorrelation between the patterns of these two conditions, particularly during the preparation and first-probe epochs. This was also the case between Prospective and Delayed trials, suggesting that mere maintenance demands did not induce marked changes in the corresponding neural codes and that it is the anticipation of interference that drives the largest changes in the representations. Importantly, we found these relationships between different conditions to be dynamic and sensitive to specific time periods within a trial. For instance, while Prospective and Delayed conditions induce the aforementioned anticorrelation during the first probe, this particular coding scheme is not observed during the second probe, where task demands are identical in these two conditions. Moreover, we observed that the transformations driven by different priorities also unfold dynamically over time. Our results showed that the neural geometry could shift from being dominated by the S-R category to being structured by the priority state (or vice versa) depending on whether the relevant content was being maintained, shielded, or executed. These results resonate with recent evidence that novel instructions recruit abstract neural codes that evolve over time depending on the specific demands of the trial epoch (Pena et al., 2025). Our findings extend these results by showing that priority demands are a crucial component with a direct impact on the representation of novel actions in WM. Perhaps more importantly, we provide evidence that the temporal dimension is relevant when assessing the relationship between active and latent representations. Our results point to a dynamic interplay between these two states, inducing important shifts to the underlying neural representations in the within-trial temporal scale.

Why might the brain implement these dynamic codes, and what computational advantages do they afford? One possibility is that the observed shifts in representational geometry may reduce potential interference. The anticipatory emergence of suppressive codes during Prospective trials suggests that the brain could actively recode WM contents in a format less vulnerable to inference when immediate execution of those contents is not required and, importantly, it is needed to perform another task. This aligns well with the idea of a protection-oriented reconfiguration, where the latent representations are reformatted into a state that minimizes the overlap with currently relevant tasks, thus reducing potential cross-talk (Stokes et al., 2020). Thus, the potential coding scheme between Current and Prospective priorities depends to an important extent on the demands of the interfering task. In our experiment, while this task (i.e., mug trials) did not share the same response mappings, its overall structure (i.e., an S-R choice reaction task) highly resembled the relevant one. This raises the possibility that the observed suppression stemmed from the high similarity between relevant and irrelevant tasks. Future studies should explore how manipulating the demands of the secondary task impacts the coding schemes of active vs. latent representations in WM.

A surprising finding in our study was the lack of evidence for S-R coding during the cue-target interval, regardless of their priority status. Instead, our analyses revealed stronger effects related to priority itself. This suggests that, while participants retained information about which mapping would be needed, the precise representational format of the S-R association was not detectable by our analytical approach. Several alternative explanations can account for this result. Primarily, it is possible that our task structure favored the storage of this information in a latent format until reactivated, as suggested by “activity-silent” proposals (Barbosa et al., 2020; Stokes, 2015). Alternatively, S-R information during the delay might involve specific neural substrates that are harder to detect with EEG than with other techniques, such as fMRI, where delay activity is frequently reported (González-García et al., 2017, 2021). This speaks to a more general limitation of our approach. While our methods are sensitive to correlations or anticorrelations across conditions, they may miss alternative, more nuanced, representational organizations. For instance, one option is that different brain regions support contents of different priority, regardless of whether they use correlated or orthogonal codes (Stokes et al., 2020). Similarly, while technically we could obtain evidence for orthogonal coding (i.e., preservation index = 0), such an estimation can be affected not only by relevant task signals, but also by unrelated noise. Future work should explore more fine-grained representational distinctions with EEG and other techniques, improving the sensitivity to latent states.

Finally, two further observations deserve attention. First, behavioral results showed faster responses in the Delayed condition compared to the Current one, which may seem counterintuitive given that the former requires maintaining S-Rs for a longer period of time. One possibility is that the retro-cue target interval may not have provided sufficient time for fully optimal anticipation, thereby hindering performance relative to Delayed trials, where participants could prepare more extensively. Second, we did not observe robust decoding of the S-R category during the initial encoding phase. This may seem surprising, given that the S-Rs were on screen during this epoch. However, similar findings have been reported in the context of novel instruction encoding, where representations of novel associations are often weak or non-discriminable at short encoding stages (Pena et al., 2025). In this regard, it is important to keep in mind that during encoding stages, multiple stimuli were simultaneously presented on screen, and participants could begin processing each piece of information differently on each trial. Such variability could negatively impact the generalizability of the EEG signal across trials. In addition, one possible explanation is that, given the novel nature of the associations on each trial, S-R representations are initially noisy or unstable and only become functionally organized later.

Altogether, our results show that WM for novel actions is an adaptive representational workspace where priority modulates not only what but also how content is maintained. While we obtained some evidence for previously proposed coding schemes (i.e., suppressive coding), our study highlights the task-optimized, dynamic nature of such strategies. As such, we observed that different conditions can relate through different coding schemes throughout a trial, depending on the specific goals and control demands.

## CRediT author contribution

J.L.D. and C.G.G. were responsible for conceptualization, project administration, and resources. C.G.G. acquired funding. I.P. performed data collection, data curation, and formal analysis. All authors were involved in methodology, visualization, and writing the original draft, reviewing, and editing.

## Data and code availability

All data and code derived from this study are available at https://osf.io/5fg7a/.

## Conflict of interest statement

The authors declare no competing financial interests.

### Acknowledgments

I.P. and C.G.G. were supported by Project PID2020-116342GA-I00 funded by MCIN/AEI/10.13039/501100011033, and Grant RYC2021-033536-I funded by MCIN/AEI/10.13039/501100011033 and by the European Union NextGeneration EU/PRTR awarded to C.G.G. J.L.D. was supported by Project PID2023-151104NA- I00 funded by MCIN/AEI/10.13039/501100011033 and by FEDER, EU, and Grant RYC2021-033940-I funded by MCIN/AEI/10.13039/501100011033 and by the European Union NextGeneration EU/PRTR. The Mind, Brain and Behavior Research Center receives funding from grants CEX2023-001312-M by MICIU/AEI/10.13039/501100011033 and UCE-PP2023-11 by the University of Granada. We thank Águeda Fuentes-Guerra and Pedro Talavera for their assistance in data collection.

## REFERENCES

1. Baddeley, A. D. (1992). Working memory. Science, 255(5044), 556–559. 10.1126/science.1736359

2. Barbosa, J., Stein, H., Martinez, R. L., Galan-Gadea, A., Li, S., Dalmau, J., Adam, K. C. S., Valls-Solé, J., Constantinidis, C., & Compte, A. (2020). Interplay between persistent activity and activity-silent dynamics in the prefrontal cortex underlies serial biases in working memory. Nature Neuroscience, 23(8), Article 8. 10.1038/s41593-020-0644-4

3. Barrouillet, P., Corbin, L., Dagry, I., & Camos, V. (2015). An empirical test of the independence between declarative and procedural working memory in Oberauer’s (2009) theory. Psychonomic Bulletin & Review, 22(4), 1035–1040. 10.3758/s13423-014-0787-y

4. Beukers, A. O., Buschman, T. J., Cohen, J. D., & Norman, K. A. (2021). Is Activity Silent Working Memory Simply Episodic Memory? Trends in Cognitive Sciences, 25(4), 284–293. 10.1016/j.tics.2021.01.003

5. Brady, T. F., Konkle, T., Alvarez, G. A., & Oliva, A. (2008). Visual long-term memory has a massive storage capacity for object details. Proceedings of the National Academy of Sciences, 105(38), 14325–14329. 10.1073/pnas.0803390105

6. Brady, T. F., Konkle, T., Alvarez, G. A., & Oliva, A. (2013). Real-world objects are not represented as bound units: Independent forgetting of different object details from visual memory. Journal of Experimental Psychology: General, 142(3), 791. 10.1037/a0029649

7. Brodeur, M. B., Guérard, K., & Bouras, M. (2014). Bank of Standardized Stimuli (BOSS) phase ii: 930 new normative photos. PLoS ONE, 9(9), e106953. 10.1371/journal.pone.0106953

8. Chun, M. M. (2011). Visual working memory as visual attention sustained internally over time. Neuropsychologia, 49(6), 1407–1409. 10.1016/j.neuropsychologia.2011.01.029

9. Cowan, N. (2010). The magical mystery four: How is working memory capacity limited, and why? Current Directions in Psychological Science, 19(1), 51–57. Scopus. 10.1177/0963721409359277

10. de Vries, I. E. J., van Driel, J., Karacaoglu, M., & Olivers, C. N. L. (2018). Priority Switches in Visual Working Memory are Supported by Frontal Delta and Posterior Alpha Interactions. Cerebral Cortex, 28(11), 4090–4104. 10.1093/cercor/bhy223

11. Formica, S., Palenciano, A. F., Vermeylen, L., Myers, N. E., Brass, M., & González- García, C. (2024). Internal attention modulates the functional state of novel stimulus-response associations in working memory. Cognition, 245, 105739. 10.1016/j.cognition.2024.105739

12. Fukuda, K., Awh, E., & Vogel, E. K. (2010). Discrete capacity limits in visual working memory. Current Opinion in Neurobiology, 20(2), 177–182. Scopus. 10.1016/j.conb.2010.03.005

13. Gade, M., Druey, M. D., Souza, A. S., & Oberauer, K. (2014). Interference within and between declarative and procedural representations in working memory. Journal of Memory and Language, 76, 174–194. 10.1016/j.jml.2014.07.002

14. González-García, C., Arco, J. E., Palenciano, A. F., Ramírez, J., & Ruz, M. (2017). Encoding, preparation and implementation of novel complex verbal instructions. NeuroImage, 148, 264–273. 10.1016/j.neuroimage.2017.01.037

15. González-García, C., Formica, S., Wisniewski, D., & Brass, M. (2021). Frontoparietal action-oriented codes support novel instruction implementation. NeuroImage, 226, 117608. 10.1016/j.neuroimage.2020.117608

16. Griffin, G., Holub, A., & Perona, P. (2006). Caltech-256 object category dataset. In *Caltech Technical Report*. 10.1021/jp953720e

17. Grootswagers, T., Wardle, S. G., & Carlson, T. A. (2017). Decoding Dynamic Brain Patterns from Evoked Responses: A Tutorial on Multivariate Pattern Analysis Applied to Time Series Neuroimaging Data. Journal of Cognitive Neuroscience, 29(4), 677–697. 10.1162/jocn_a_01068

18. Guggenmos, M., Sterzer, P., & Cichy, R. M. (2018). Multivariate pattern analysis for MEG: A comparison of dissimilarity measures. NeuroImage, 173, 434–447. 10.1016/J.NEUROIMAGE.2018.02.044

19. Konkle, T., Brady, T. F., Alvarez, G. A., & Oliva, A. (2010). Conceptual distinctiveness supports detailed visual long-term memory for real-world objects. Journal of Experimental Psychology: General, 139(3), 558. 10.1037/a0019165

20. Kriegeskorte, N., Mur, M., & Bandettini, P. (2008). Representational similarity analysis—Connecting the branches of systems neuroscience. Frontiers in Systems Neuroscience, 2(November), 4. 10.3389/neuro.06.004.2008

21. Linde-Domingo, J., & Spitzer, B. (2024). Geometry of visuospatial working memory information in miniature gaze patterns. Nature Human Behaviour, 8(2), 336–348. 10.1038/s41562-023-01737-z

22. Meiran, N., Pereg, M., Kessler, Y., Cole, M. W., & Braver, T. S. (2015). The power of instructions: Proactive configuration of stimulus–response translation. *Journal of Experimental Psychology: Learning*, Memory, and Cognition, 41(3), 768– 786. 10.1037/xlm0000063

23. Muhle-Karbe, P. S., Myers, N. E., & Stokes, M. G. (2021). A hierarchy of functional states in working memory. Journal of Neuroscience, 41(20), 4461–4475. 10.1523/JNEUROSCI.3104-20.2021

24. Myers, N. E., Chekroud, S. R., Stokes, M. G., & Nobre, A. C. (2018). Benefits of flexible prioritization in working memory can arise without costs. Journal of Experimental Psychology. Human Perception and Performance, 44(3), 398–411. 10.1037/xhp0000449

25. Myers, N. E., Stokes, M. G., & Nobre, A. C. (2017). Prioritizing Information during Working Memory: Beyond Sustained Internal Attention. Trends in Cognitive Sciences, 21(6), 449–461. 10.1016/j.tics.2017.03.010

26. Nobre, A. C. (Kia), & Stokes, M. S. (2020). Memory and Attention: The Back and Forth. In D. Poeppel, G. R. Mangun, & M. S. Gazzaniga (Eds.), *The Cognitive Neurosciences* (p. 0). The MIT Press. 10.7551/mitpress/11442.003.0035

27. Oberauer, K. (2009). Design for a working memory. Psychology of Learning and Motivation, 51, 45–100.

28. Oberauer, K. (2010). Declarative and Procedural Working Memory: Common Principles, Common Capacity Limits? Psychologica Belgica, 50(3–4). 10.5334/pb-50-3-4-277

29. Olmos-Solis, K., van Loon, A. M., & Olivers, C. N. L. (2021). Content or status: Frontal and posterior cortical representations of object category and upcoming task goals in working memory. Cortex, 135, 61–77. 10.1016/j.cortex.2020.11.011

30. Peirce, J. W. (2007). PsychoPy—Psychophysics software in Python. Journal of Neuroscience Methods, 162(1), 8–13. 10.1016/j.jneumeth.2006.11.017

31. Pena, P., Palenciano, A. F., González-García, C., & Ruz, M. (2025). Novel Verbal Instructions Recruit Abstract Neural Patterns of Time-Variable Information Dimensionality. Journal of Neuroscience, 45(17). 10.1523/JNEUROSCI.1964-24.2025

32. Schneegans, S., & Bays, P. M. (2017). Neural Architecture for Feature Binding in Visual Working Memory. Journal of Neuroscience, 37(14), 3913–3925. 10.1523/JNEUROSCI.3493-16.2017

33. Shepherdson, P., Oberauer, K., & Souza, A. S. (2018). Working memory load and the retro-cue effect: A diffusion model account. Journal of Experimental Psychology: Human Perception and Performance, 44(2), 286–310. 10.1037/xhp0000448

34. Stokes, M. G. (2015). “Activity-silent” working memory in prefrontal cortex: A dynamic coding framework. Trends in Cognitive Sciences, 19(7), 394–405. 10.1016/j.tics.2015.05.004

35. Stokes, M. G., Muhle-Karbe, P. S., & Myers, N. E. (2020). Theoretical distinction between functional states in working memory and their corresponding neural states. Visual Cognition, 28(5–8), 420–432. 10.1080/13506285.2020.1825141

36. van Ede, F. (2020). Visual working memory and action: Functional links and bi- directional influences. Visual Cognition, 1–13. 10.1080/13506285.2020.1759744

37. van Loon, A. M., Olmos-Solis, K., Fahrenfort, J. J., & Olivers, C. N. (2018). Current and future goals are represented in opposite patterns in object-selective cortex. eLife, 7, e38677. 10.7554/eLife.38677

38. Walther, A., Nili, H., Ejaz, N., Alink, A., Kriegeskorte, N., & Diedrichsen, J. (2016). Reliability of dissimilarity measures for multi-voxel pattern analysis. NeuroImage, 137, 188–200. 10.1016/J.NEUROIMAGE.2015.12.012

39. Wan, Q., Menendez, J. A., & Postle, B. R. (2022). Priority-based transformations of stimulus representation in visual working memory. PLOS Computational Biology, 18(6), e1009062–e1009062. 10.1371/journal.pcbi.1009062

40. Yu, Q., Teng, C., & Postle, B. R. (2020). Different states of priority recruit different neural representations in visual working memory. PLOS Biology, 18(6), e3000769. 10.1371/journal.pbio.3000769

